# ILC1-derived IFN-γ mediates cDC1-dependent host resistance against *Toxoplasma gondii*

**DOI:** 10.1101/2020.01.06.895771

**Authors:** Américo H. López-Yglesias, Elise Burger, Ellie Camanzo, Andrew T. Martin, Alessandra M. Araujo, Samantha F. Kwok, Felix Yarovinsky

## Abstract

Host resistance against intracellular pathogens requires a rapid IFN-γ mediated immune response. We reveal that T-bet-dependent production of IFN-γ is essential for the maintenance of inflammatory DCs at the site of infection with a common protozoan parasite, *Toxoplasma gondii*. A detailed analysis of the cellular sources for T-bet-dependent IFN-γ identified that ILC1s, but not NK or T_H_1 cells, were involved in the regulation of inflammatory DCs via IFN-γ. Mechanistically, we established that T-bet dependent ILC1-derived IFN-γ is critical for the induction of IRF8, an essential transcription factor for cDC1s. Failure to upregulate IRF8 in DCs resulted in acute susceptibility to *T. gondii* infection. Our data identifies that ILC1-derived IFN-γ is indispensable for host resistance against intracellular infection via maintaining IRF8+ inflammatory DCs at the site of infection.

**Author Summary:** Mounting a robust type I innate immune response is essential for resistance against numerous intracellular pathogens. The type I immune response is characterized by the production of IFN-γ, a central cytokine required for multiple non-redundant effector functions against bacterial, viral, and parasitic pathogens. Previous work has shown that group 1 innate lymphocyte cells (ILC1s) together with NK and CD4+ T cells play an indispensable IFN-γ mediated protective role against *Toxoplasma gondii* infection; yet, the pathway of how IFN-γ produced by ILC1s defend against *T. gondii* remains unknown. In this work we identified that early production of IFN-γ by ILC1 is essential for maintaining dendritic cells (DCs) during infection. Mechanistically, we reveal that ILC1-derived IFN-γ is indispensable for inducing the transcription factor IRF8 that is critical for sustaining inflammatory DCs. Finally, we demonstrate that IRF8+ DCs are critical for parasite elimination.

## INTRODUCTION

Host defense against intracellular pathogens requires a quick and effective type I immune response. A coordinated response of innate myeloid and lymphoid cells is critical for both rapid pathogen restriction and activation of the adaptive immune response. The transcription factor T-bet, encoded by *Tbx21*, has been shown to play a critical role for the effector function of innate and adaptive lymphocytes in response to intracellular pathogens via regulation of IFN-γ production [1–8]. The cytokine IFN-γ is indispensable for host defense as it is essential for the induction of anti-microbial IFN-γ-inducible genes, which results in intracellular microbial clearance.

The obligate intracellular protozoan parasite, *Toxoplasma gondii*, is a potent inducer of IFN-γ and has been exploited to characterize the host’s innate and adaptive type I immune responses against intracellular pathogens [9]. Immunity against *T*. *gondii* requires a type I CD4+ T helper cell (T_H_1)-derived IFN-γ response, and in the absence of either CD4+ T cells or IFN-γ, the host rapidly succumbs to infection [10–12]. Therefore, it was anticipated that the rapid susceptibility observed in *T*. *gondii* infected T-bet-deficient (*Tbx21*^−/−^) mice was due to the absence of CD4+ T cell-derived IFN-γ. However, our group and others have recently observed that *Tbx21*^−/−^ mice maintained IFN-γ producing CD4+ T cells during parasite infection, while remaining highly susceptible to infection [4, 5]. These data suggest an innate T-bet-dependent mechanism that is critical for a protective type I immune response against *T*. *gondii* infection.

Natural killer (NK) cells and group 1 innate lymphoid cells (ILC1s) are critical sources of innate IFN-γ during intracellular infection [13, 14]. It has been established that the transcription factors, Eomesodermin (Eomes) and T-bet play a role in NK cell development [15, 16]. However, while Eomes is indispensable for NK cell maturation and effector function, T-bet plays a more limited role in the development, migration, and cytokine production of NK cells [4, 16–18]. During *T*. *gondii* infection, NK cell-derived IFN-γ stimulates the effector function of inflammatory myeloid cells, and in the absence of NK cells the host immunity to *T*. *gondii* is compromised [13, 19].

Similar to NK cells and T_H_1s, tissue resident ILC1s have been identified as a critical source of IFN-γ necessary for pathogen restriction [14, 20–22]. Unlike NK cells, the maturation and cytokine production of ILC1s is T-bet-dependent and Eomes-independent [14]. Tissue resident ILC1s rapidly respond to type I conventional DC (cDC1)-derived IL-12 in an antigen independent manner, leading to IFN-γ production [22, 23]. In addition to providing an early source of IFN-γ, ILC1s can also augment the recruitment of innate inflammatory myeloid cells during *T*. *gondii* infection [14], demonstrating that crosstalk between innate myeloid cells and ILC1s is a critical component of an effective innate type I immune response against intracellular pathogens.

Because multiple cellular sources of IFN-γ play a complex and non-redundant role for T-bet-dependent host defense against a common protozoan parasite *T. gondii*, we sought to determine the role T-bet plays in coordinating innate myeloid cells and lymphocytes to work in concert with one another for host resistance. Our experiments revealed that in the absence of T-bet, inflammatory DCs were significantly compromised at the site of infection. We identified that parasite-mediated T-bet-dependent ILC1-derived IFN-γ is crucial for maintaining inflammatory DCs during infection. Importantly, the absence of DCs could be rescued by exogenous administration of IFN-γ during *T*. *gondii* infection, indicating a critical role for early T-bet-dependent IFN-γ in regulating inflammatory DCs. Mechanistically, we uncovered that ILC1-derived IFN-γ was required for induction of the transcription factor interferon regulatory factor-8 (IRF8) in cDC1s, which are required for immunity against *T. gondii*. Our results establish that during *T. gondii* infection, T-bet-dependent ILC1-derived IFN-γ is indispensable for regulating inflammatory IRF8+ cDC1s, leading to parasite clearance and host survival.

## RESULTS

### T-bet is critical to maintain inflammatory DCs during *T. gondii* infection

To establish T-bet’s protective role in host defense against intracellular pathogens, we implemented the well-established intraperitoneal (i.p.) model of infection with *T*. *gondii* that triggers and depends on a robust CD4+ T_H_1-derived IFN-γ response [10, 24]. Considering the importance of T-bet in the regulation of T_H_1 cells [7], we anticipated that T-bet-deficient mice would demonstrate enhanced susceptibility to *T*. *gondii* similar to T-cell deficient mice due to the lack of T-bet-mediated T_H_1-derived IFN-γ production. Strikingly, mice lacking T-bet were more susceptible to *T. gondii* compared to *RAG2*^−/−^ animals and were unable to survive past day 10 of the acute stage of infection (Fig. 1A). These data suggested that the transcription factor T-bet is involved in innate host defense and essential for host survival through the acute stage of parasitic infection independently of the regulation of T_H_1 immunity to the parasite.

**Figure 1.**
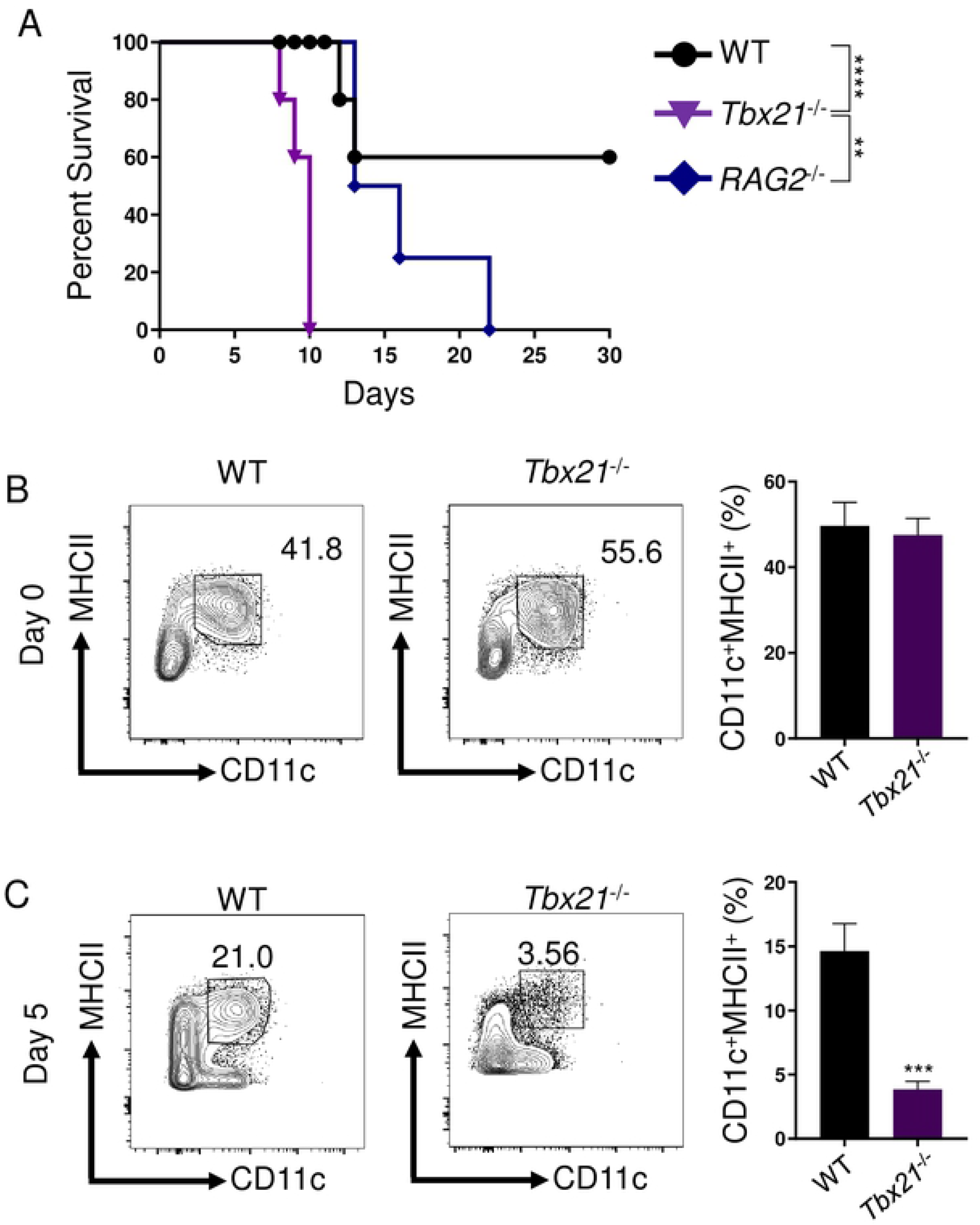
*T-bet is critical for inflammatory DCs and host resistance during acute* T. gondii *infection*. (**A**) Survival of WT (●), *Tbx21*^−/−^ (▼), and *RAG2*^−/−^ (♦) mice infected i.p. with 20 cysts of the ME49 strain of *T*. *gondii*. WT and *Tbx21*^−/−^ mice were infected i.p. with 20 cysts of *T*. *gondii*. PECs were harvested on days 0 and 5 (**B, C**) and Lin-CD11c+MHCII+ DCs and were analyzed by flow cytometry. Average frequencies of Lin-CD11c+MHCII+ DCs (B, C) in the PECs were analyzed on days 0 and 5 following infection. Results are representative of three-independent experiments involving at least 3 mice per group. Statistical analyses were done using Log-rank (Mantel Cox) test or unpaired t-test analysis of individual groups, \*\**P*<0.01, \*\*\**P*<0.001, \*\*\*\**P*<0.0001. Error bars, standard error mean.

It has been well-established that *T*. *gondii*-mediated type I immunity requires DCs for host defense [25–29]. Work from our lab and others have demonstrated that DC-deficiency results in acute host susceptibility to *T*. *gondii* [26, 30]. Therefore, we examined if the lack of T-bet compromised inflammatory DC-mediated immunity. We observed that in naïve mice, lack of T-bet had no discernable effects on the presence of DCs in the peritoneal cavity (Fig. 1B), defined as Lin^−^CD11c^+^MHCII^+^ cells (Fig. S5). In striking contrast, *Tbx21*^−/−^ mice had significantly reduced inflammatory DCs compared to both WT and *RAG2*^−/−^ animals when infected with *T. gondii* (Fig. 1C, Fig. S1C), independent of intrinsic T-bet expression by Lin^−^CD11c^+^MHCII^+^ DCs (data not shown). A detailed analysis of inflammatory DCs present at the site of the infection revealed that as early as day 3 post-infection, there was a noticeable reduction in DCs analyzed in *Tbx21*^−/−^ mice when compared to WT mice (Fig. S1A). Furthermore, by day 5 post-infection, inflammatory DCs were practically undetectable in the absence of T-bet, a trend that was also observed by day 8 following infection (Fig. 1C, Fig. S1B). Overall, our experiments established a critical innate function of T-bet for regulating the presence of inflammatory DCs during acute parasitic infection.

### Early T-bet-dependent IFN-γ is critical for inflammatory DCs via regulation of IRF8

The IFN-γ inducible transcription factor IRF8, originally described as interferon consensus sequence-binding protein (ICSBP), is known to play a key role in the development and survival of cDC1s, also known as CD8^+^BATF3^+^ DCs [31–35]. Therefore, we investigated if T-bet regulates the presence of inflammatory DCs via induction of IRF8. We observed that in naïve mice, T-bet was dispensable for the presence of a small population of peritoneal IRF8+ DCs, similar to WT and *RAG2*^−/−^ animals (Fig. 2A, B & Fig. S1D). Strikingly, *T*. *gondii* infection resulted in the rapid accumulation of inflammatory IRF8+ DCs in WT mice and this population was virtually absent in *Tbx21*^−/−^ mice by day 8 post-infection (Fig. 2C-F). These data suggest that early IFN-γ missing in *Tbx21*^−/−^ mice results in the absence of cDC1s.

**Figure 2.**
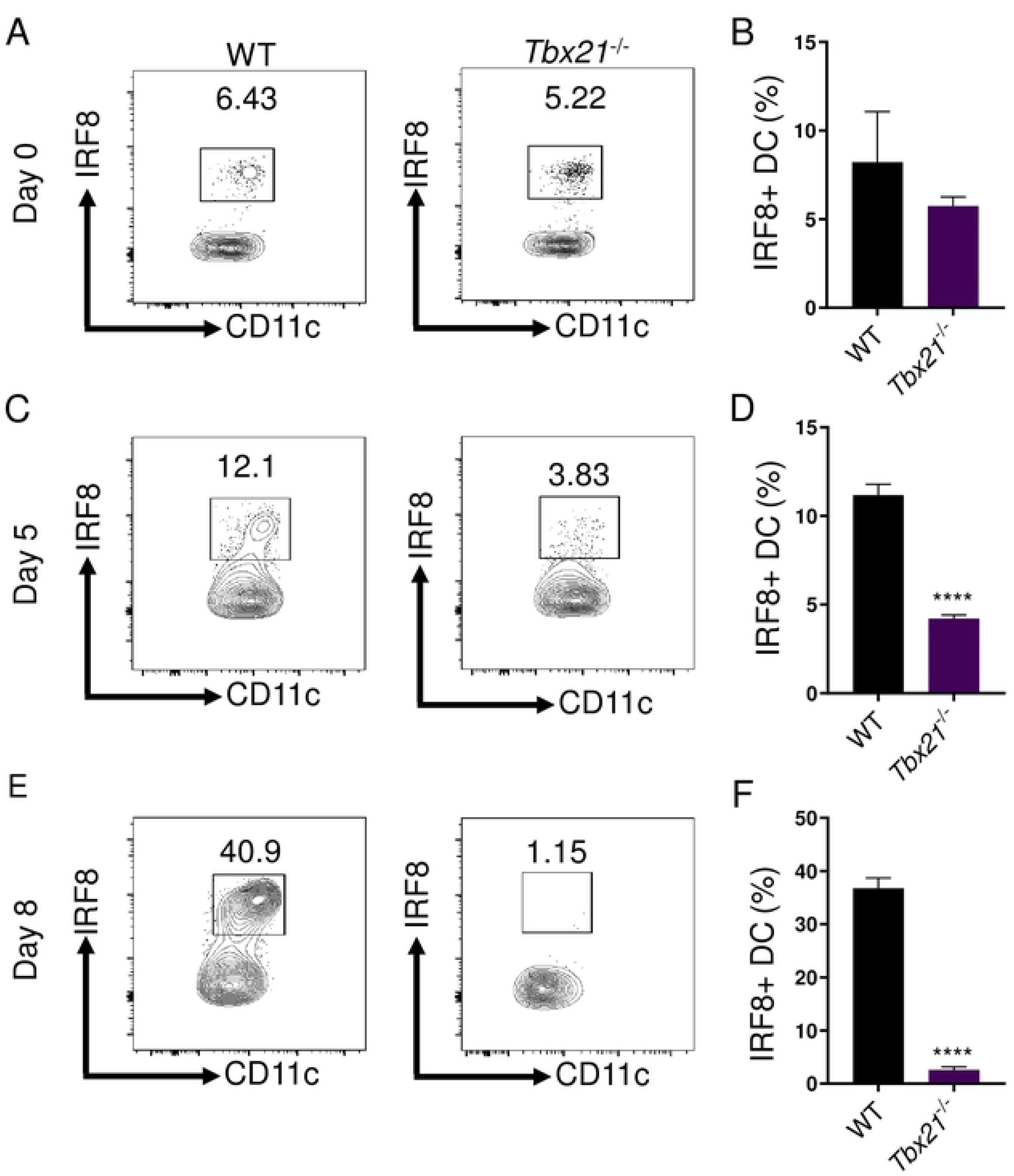
T-bet is essential for sustaining IRF8+ DCs. (**A-F**) WT and *Tbx21*^−/−^ mice were infected i.p. with 20 cysts of *T*. *gondii*. (**B**, **D**, **F**) Average frequencies of Lin- CD11c+MHCII+IRF8+ DCs in the PECs were analyzed on days 0, 5, and 8 following infection of A, C, E. Results are representative of three-independent experiments involving at least 3 mice per group. Statistical analyses were done using unpaired t-test analysis of individual groups, \*\*\*\**P*<0.0001. Error bars, standard error mean.

To identify if the lack of T-bet resulted in impaired early IFN-γ production during *T*. *gondii* infection we performed a detailed analysis of *T*. *gondii*-mediated IFN-γ production. Our results identified that T-bet was critical for early IFN-γ production on days 3 and 5 of parasite infection (Fig. 3A). Therefore, we hypothesized early IFN-γ is required to maintain DCs during *T*. *gondii* infection. To test our hypothesis, IFN-γ was administered to infected *Tbx21*^−/−^ mice, which not only significantly augmented inflammatory DCs at the site of infection (Fig. 3B, C), but also rescued IRF8 expression in DCs (Fig. 3D-F). These data demonstrate T-bet-dependent early IFN-γ production is essential for maintaining inflammatory DCs via regulation of IRF8.

**Figure 3.**
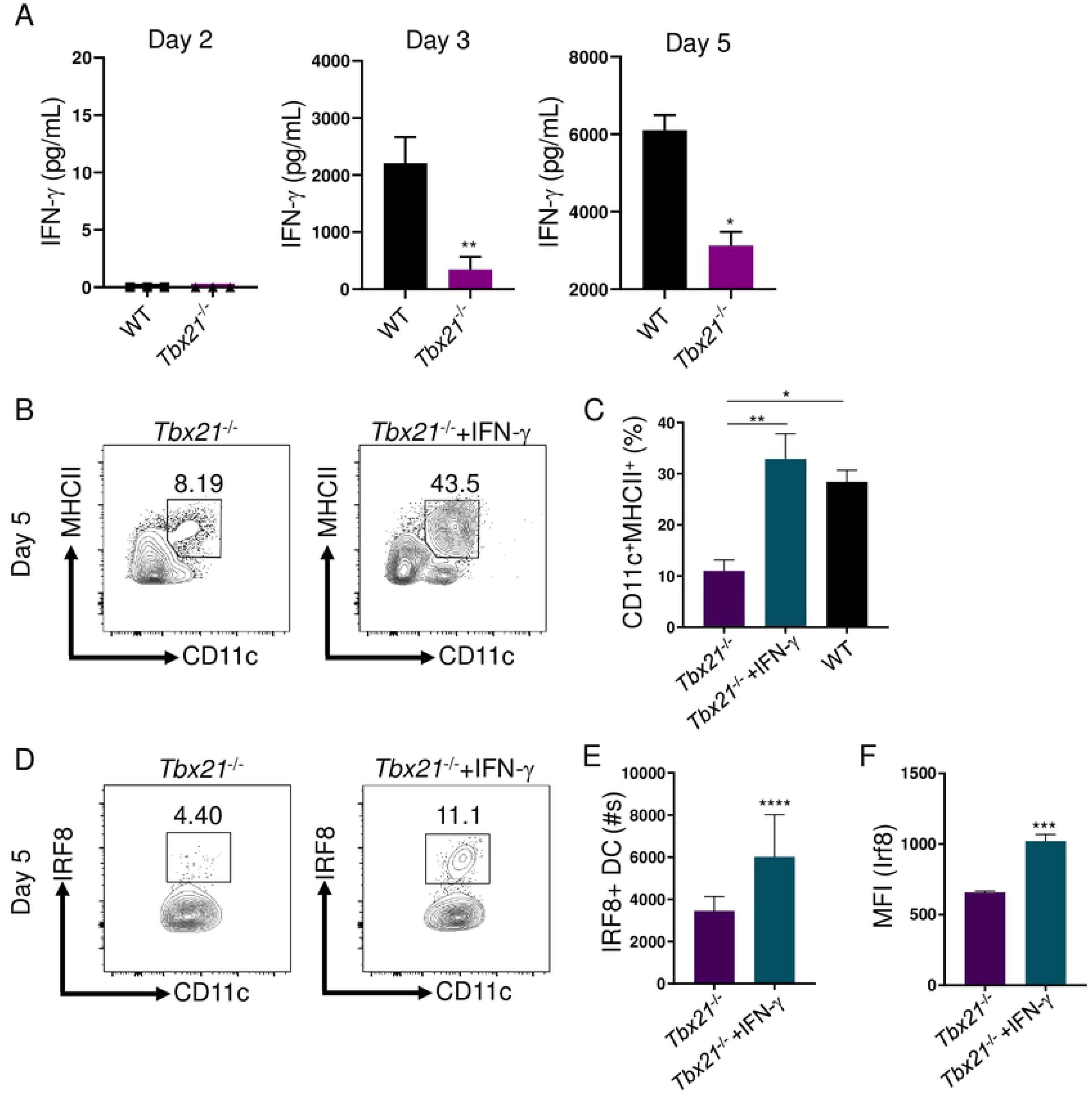
Early T-bet-dependent IFN-γ is critical to maintain inflammatory IRF8+ cDCs1. (**A**) IFN-γ analysis by ELISA of serum in mice following *T*. *gondii* infection on days 2, 3, and 5 post-infection. (**B-F**) *Tbx21*^−/−^ mice were i.p. infected with *T*. *gondii* and treated with or without IFN-γ. (C) Average frequencies of Lin-CD11c+MHCII+ DCs in the PECs were analyzed on day 5 following infection. Absolute quantification of (E) Lin-CD11c+MHCII+IRF8+ DCs in the PECs were analyzed on day 5 following infection. (F) Mean fluorescent intensity (MFI) of Lin- CD11c+MHCII+ DC IRF8 expression in the PECs was analyzed on day 5 post-infection. Results are representative of three-independent experiments involving at least 3 mice per group. Statistical analyses were done using (A, E, F) unpaired t-test analysis of individual groups or (C) one-way Anova with Tukey’s multiple comparison test, \**P*<0.05, \*\**P*<0.01, \*\*\**P*<0.001, \*\*\*\**P*<0.0001. Error bars, standard error mean.

### T-bet regulates ILC1-derived IFN-γ, but is dispensable for NK cell-derived IFN-γ

To thoroughly define the function of IFN-γ in regulating inflammatory DCs, we examined if neutralizing IFN-γ eliminates their presence at the site of infection. We observed that blocking IFN-γ during *T*. *gondii* infection resulted in a striking reduction of inflammatory DCs in both WT and T cell-deficient mice (Fig. 4A-C). These results revealed that IFN-γ from innate immune cells was sufficient to maintain inflammatory DCs in the absence of T cell-derived IFN-γ. Among innate immune functions, T-bet has been described to play an important role for NK cell function and ILC1 development, and both of these cell types primarily control acute *T*. *gondii* infection via their production of IFN-γ [14–16]. Therefore, we examined the importance of innate IFN-γ for the maintenance of inflammatory DCs. Treatment of WT and *RAG2*^−/−^ mice with anti-NK1.1 antibody resulted in a profound loss of inflammatory DCs (Fig. 4A-C), further identifying that innate lymphoid cells are critically involved in sustaining parasite-triggered inflammatory DCs. To further explore if ILC1s were sufficient and necessary for maintaining DCs during infection, we next explored the effects of selective depletion of Thy1+ cells in WT and *RAG2*^−/−^ mice. Depletion of Thy1 expressing cells resulted in a significant decrease of DCs in WT and *RAG2*^−/−^ mice (Fig. 4A-D). Moreover, Thy1-mediated depletion of ILC1s in *RAG2*^−/−^ mice resulted in an increase of pathogen burden to the levels seen in T-bet-deficient animals (Fig. S2). Our data implicates that either NK cell- or ILC1-derived IFN-γ, in a T-bet-dependent manner, are required to maintain inflammatory DCs during *T*. *gondii* infection.

**Figure 4.**
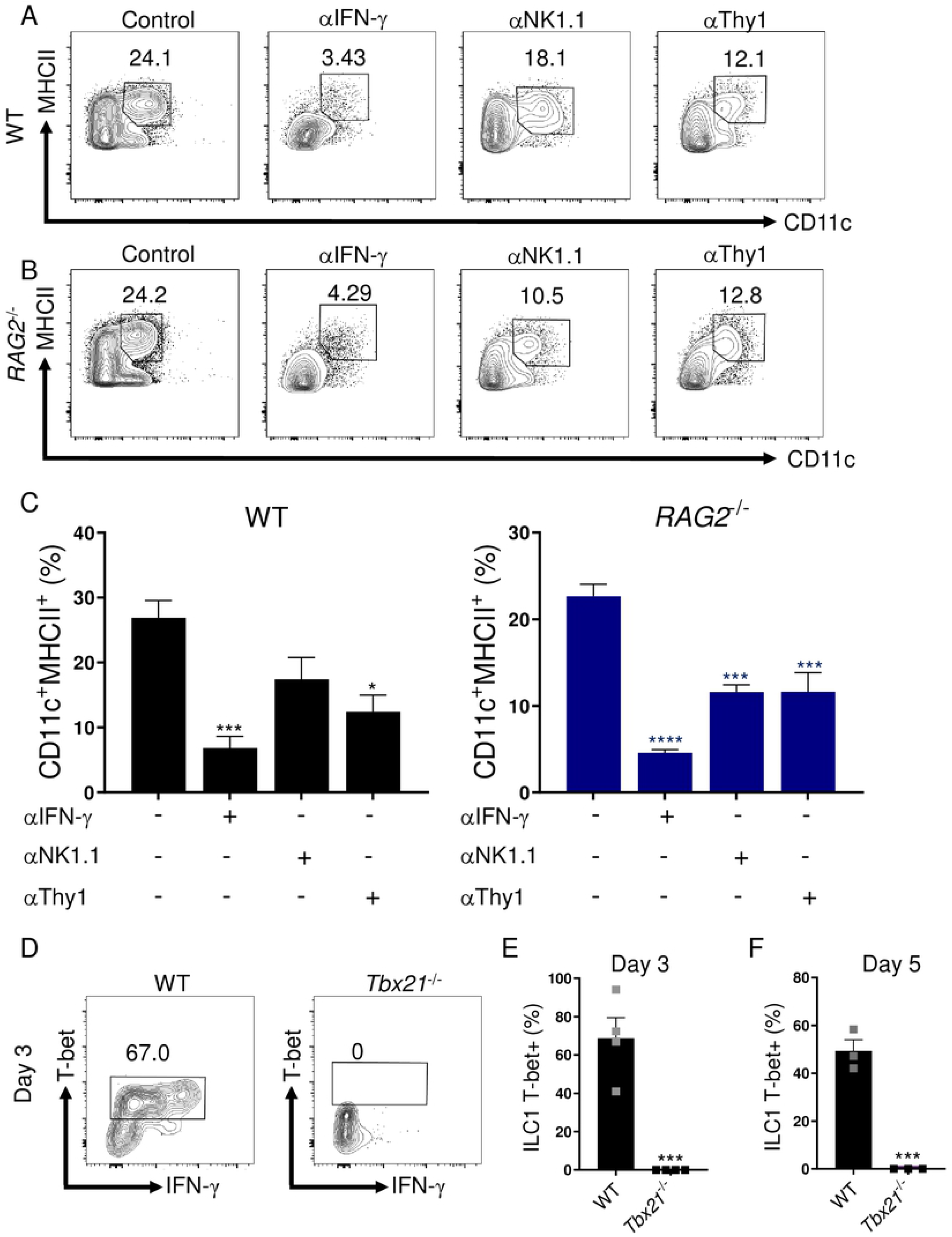
*IFN-γ and ILC1s are required for sustaining inflammatory DCs during* T. gondii *infection*. (**A, B**) WT and *RAG2*^−/−^ mice were i.p. infected with *T*. *gondii* and then treated with anti-IFN-γ, anti-NK1.1, or anti-Thy1 antibodies during infection. (**C**) Average frequencies of CD11c+MHCII+ DCs in the PECs were analyzed on day 8 following infection and antibody treatment. (**D-F**) WT and *Tbx21*^−/−^ mice were infected i.p. with 20 cysts of *T*. *gondii*. (D-F) Average frequencies of CD127+NKp46+T-bet+ ILC1s cells in the PECs were analyzed on days 3 and 5 following infection. Results are representative of three-independent experiments involving at least 3 mice per group. Statistical analyses were done using (E, F) unpaired t-test analysis of individual groups or (C) one-way Anova with Tukey’s multiple comparison test, \**P*<0.05, \*\*\**P*<0.001, \*\*\*\**P*<0.0001. Error bars, standard error mean.

Our experiments with WT and T-bet-deficient mice revealed that in response to *T. gondii* infection, NK cells produced large amounts of IFN-γ in a T-bet-independent manner. This was evident from comparable production of IFN-γ by NK cells seen in WT and *Tbx21*^−/−^ mice on days 3 and 5 post-infection (Fig. S3). Thus, T-bet is not required for IFN-γ production by NK cells during *T*. *gondii* infection, suggesting that T-bet-dependent regulation of inflammatory DCs is not mediated by NK-derived IFN-γ.

As expected, we observed that WT mice retained robust T-bet expressing ILC1-derived IFN-γ responses during infection (Fig. 4E-G). We identified ILC1s from the peritoneal cavity as CD3^−^CD19^−^Ly6G^−^NKp46^+^CD127^+^ (Fig. S5) and upon further evaluation, confirmed this population was also GranzymeB^−^CD49b^−^CD49a^+^CD200R^+^ (Fig. S4), and as anticipated were absent in *Tbx21*^−/−^ mice at all examined time points following *T*. *gondii* infection (Fig. 4E-G). Therefore, T-bet deficiency resulted in two major defects in innate immunity: lack of ILC1s and inflammatory DCs at the site of infection. We further examined a role for ILC1 in maintenance of inflammatory DCs during *T*. *gondii* infection by analyzing parasite infected lymphocyte-deficient (*RAG2*^−/−^γc^−/−^) mice, which also lack ILC1s. We observed that similar to T-bet deficiency, *RAG2*^−/−^γc^−/−^ mice had very few inflammatory DCs (data not shown), further demonstrating that ILC1s are essential for the presence of inflammatory DCs during parasite infection.

### IRF8+ DCs are critical for pathogen clearance against intracellular infection

Our data has identified that early ILC1-derived IFN-γ is critical for maintaining IRF8+ DCs during infection and limiting parasite replication throughout the host. Therefore, we hypothesized that conditionally deleting IRF8 expression by DCs would result in uncontrolled parasite replication during infection and rapid host mortality. To test our hypothesis, we infected mice with a DC-restricted deficiency of IRF8 by using the CD11c-Cre system (CD11c-Cre x *Irf8*^flox/flox^ mice, DC-*Irf8*^−/−^). Infected DC-*Irf8*^−/−^ mice revealed an overall reduction of total MHCII^+^CD11c^+^ DCs by day 5 post-infection (Fig. 5A). Moreover, the absence of IRF8 expression in MHCII^+^CD11c^+^ DCs resulted in dramatically elevated parasite burden both at the site of infection and in the spleen on day 5 post infection (Fig. 5B, C). We then examined if IRF8-deficient DCs were required for host survival against *T*. *gondii*. Similarly to T-bet-deficient mice, DC-*Irf8*^−/−^ animals succumb to parasite infection rapidly (Fig. 5D). Our data establishes that early and rapid T-bet-dependent ILC1-derived IFN-γ is indispensable for maintaining IRF8+ cDC1s, which are required for pathogen clearance and host survival.

**Figure 5.**
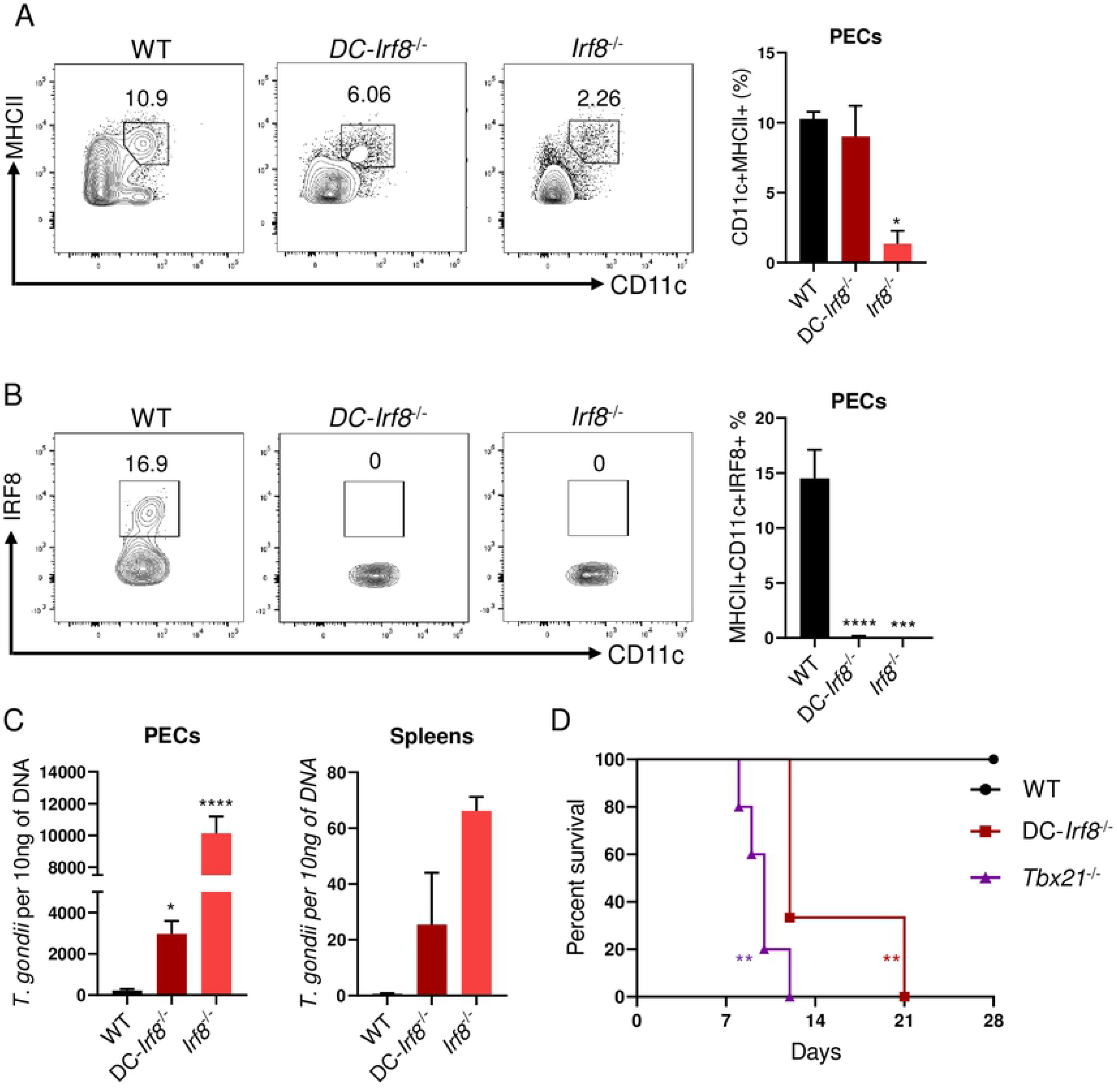
cDC1s are essential for host resistance against intracellular infection. (**A**, **B, C**) WT, DC-*Irf8*^−/−^, and *Irf8*^−/−^ mice were infected i.p. with 20 cysts of *T*. *gondii*. (A, B) Average frequencies of Lin^−^CD11c^+^MHCII^+^ and Lin^−^CD11c^+^MHCII^+^IRF8^+^ DCs in the PECs were analyzed on day 5 following infection of A, B. (C) WT, DC-*Irf8*^−/−^, and *Irf8*^−/−^ mice were i.p. infected with *T*. *gondii* and pathogen burden was assessed from PECs and spleen of mice following infection by qPCR on day 5 post-infection. (**D**) Survival of WT (●), *Tbx21*^−/−^ (▲), and DC-*Irf8*^−/−^ (■) mice infected i.p. with 20 cysts of *T*. *gondii*. Results are representative of three-independent experiments involving at least 3 mice per group. Statistical analyses were done using Log-rank (Mantel Cox) test or one-way Anova with Tukey’s multiple comparison test, \**P*<0.05, \*\*\**P*<0.001, \*\*\*\**P*<0.0001. Error bars, standard error mean.

## DISCUSSION

The cytokine IFN-γ is critical for triggering cellular anti-parasitic defense mechanisms that are essential for the destruction of the intracellular parasitophorous vacuole and *T. gondii* clearance [9, 12, 36–43]. Classically, the transcription factor T-bet is considered the master regulator for determining the CD4+ T_H_1 lineage and IFN-γ production [7]. However, recent studies revealed that that T-bet is largely dispensable for CD4+ T cell-derived IFN-γ responses [4, 5]. Herein, we demonstrate *Tbx21*^−/−^ mice succumb to infection significantly quicker than mice lacking T cells, suggesting an innate T-bet-dependent mechanism of host immunity, critical for survival of the acute stage of infection.

Innate lymphoid cells are critical early responders to intracellular pathogens. During *T*. *gondii* infection, two sets of ILCs are known to have essential roles for host resistance, NK cells, and ILC1s. T-bet is associated with the maturation, egress, and effector function of NK cells [15, 17, 18]. Yet, NK cell development is not impaired in the absence of T-bet in this model [16]. Moreover, *T*. *gondii*-triggered NK cell-derived IFN-γ production plays a critical role for initiating the effector function of myeloid cells that are required for host resistance [13, 19, 44]. Our results demonstrate that the absence of T-bet does not impede the recruitment of NK cells to the site of infection, nor does it abolish their capability to produce IFN-γ. These data establish that during *T*. *gondii* infection, T-bet plays a limited role for the migration of NK cells to the peritoneum and their IFN-γ production.

Along with NK cells, ILC1s have been shown to be a critical early source of IFN-γ during microbial invasion. Early ILC1-derived IFN-γ is critical for host resistance against Mouse Cytomegalovirus (MCMV), *Clostridium*, *Salmonella*, and *T*. *gondii* [14, 20–22]. Moreover, T-bet is required for the maturation and cytokine production of ILC1s [14]. Previous groups have observed that *T. gondii* infection mediates ILC1-derived IFN-γ and plays an important role in parasite clearance [14]. By examining *T. gondii*-infected *Tbx21*^−/−^ mice, we were able to define a host defense function of T-bet-dependent ILC1s during *T*. *gondii* infection. T-bet-deficiency resulted in the loss of ILC1s, but did not abolish IFN-γ producing NK cells or CD4+ T cells [5, 14]. This suggested that the significant reduction of inflammatory DCs observed in *Tbx21*^−/−^ mice was caused by the absence of T-bet-dependent ILC1-derived IFN-γ. By antibody depletion of ILC1s or IFN-γ neutralization, we revealed that ILC1-derived IFN-γ plays a critical role in maintaining inflammatory DCs and restricting parasite growth both locally and in peripheral tissues. Additionally, we have observed that ILC3s do not play a role in host defense against *T*. *gondii* (data not shown). Thus, we show that during *T*. *gondii* infection, the critical function of T-bet-dependent ILC1-derived IFN-γ is to maintain inflammatory DCs at the site of infection.

We and others have previously established that the transcription factor IRF8 is essential for host resistance to *T*. *gondii* [34, 45]. IRF8-deficiency resulted in acute susceptibility to *T*. *gondii* due to the absence of cDC1s [46]. Our results revealed that early T-bet-dependent IFN-γ was critical for maintaining inflammatory DCs via regulation of IRF8 expression and DC-specific IRF8 is required for host resistance to the parasite.

This study defines the critical function of early T-bet-dependent IFN-γ in sustaining inflammatory IRF8+ DCs during intracellular pathogen infection. Our results demonstrate continuous crosstalk between DCs and ILC1s mediated by IL-12 and IFN-γ, where DC-derived IL-12 triggers an early IFN-γ response from ILC1s [22] and IFN-γ produced by group 1 innate lymphoid cells is essential to maintaining inflammatory IRF8+ DCs during infection, which are indispensable for host immunity.

## MATERIAL AND METHODS

### Animals

C57BL/6, *RAG2*^−/−^, *Tbx21*^−/−^, CD11c-Cre, and *Irf8*^flox/flox^ mice were obtained from Jackson Laboratory (Bar Harbor, ME). All control and experimental mice were age- and sex-matched within all individual experiments. This study included both male and female mice, and the data derived from male and female mice identified no sex-specific differences in the performed experiments.

### Ethics Statement

All mice were maintained at in the pathogen-free American Association of Laboratory Animal Care-accredited animal facility at the University of Rochester Medical Center, Rochester, NY. All animal experimentation (animal protocol #102122) has been reviewed and approved by the University Committee on Animal Resources (UCAR). UCAR inspects, at least once every six months, all of the animal facilities, including animal study areas using the USDA Regulations and The Public Health Service (PHS), as basis

#### Toxoplasma gondii infection and qPCR

All mice were i.p. infected with an average of 20 *T*. *gondii* cysts of the ME49 strain. At days 0, 3, 5, and 8 post-infection, the animals were necropsied. In some experiments, mice were injected i.p. with 50 ng of IFN-γ (R&D Systems) on days 0, 1, 2, and 3. To determine *T*. *gondii* pathogen loads, total genomic DNA from animal tissue was isolated by using the DNeasy Blood and Tissue Kit (Qiagen) according to manufacturer’s instructions. PCR were performed by using SSOFast Eva Green Supermix (BioRad). Samples were measured by qPCR using a MyiQ Real-Time PCR Detection System (BioRad), and data from genomic DNA was compared with a defined copy number standard of the *T*. *gondii* gene *B1*.

#### ELISA Analysis

The IFN-γ concentration in the sera was analyzed by standard sandwich ELISA kit according to manufacturer’s instructions (ThermoFisher).

#### Measurements of immune cells responses

To assay the responses of mice infected with *T*. *gondii* the PECs were harvested from C57BL/6, *RAG2*^−/−^, and *Tbx21*^−/−^ mice on days 0, 3, 5, and 8 post-infection. To examine NK cells and ILC1s responses single-cell suspension of PECs were restimulated with PMA (20 ng/mL) and ionomycin (1 μg/mL) (Sigma-Aldrich) for 4 hours in the presence of GolgiPlug (Brefeldin A, BD Biosciences). After isolation and *in vitro* restimulation, the cells were washed once in phosphate-buffered saline and stained with Zombie Yellow (BioLegend) to assess live vs. dead status of cells. Cells were then washed with phosphate-buffered saline + 1% fetal bovine serum and stained with fluorochrome-conjugated antibodies. For intracellular staining and subsequent washing, cells were permeabilized overnight at 4°C with the Foxp3/ Transcription Factor Staining Buffer Set according to the manufacturer’s instructions (ThermoFisher).Cell fluorescence was measured using an LSRII flow cytometer, and data were analyzed using FlowJo Software (Tree Star, Ashland, OR).

### Statistical Analysis

All data were analyzed with Prism (Version 8; GraphPad, La Jolla, CA). These data were considered statistically significant when *P*-values were <0.05.

---

**Figure S1.**
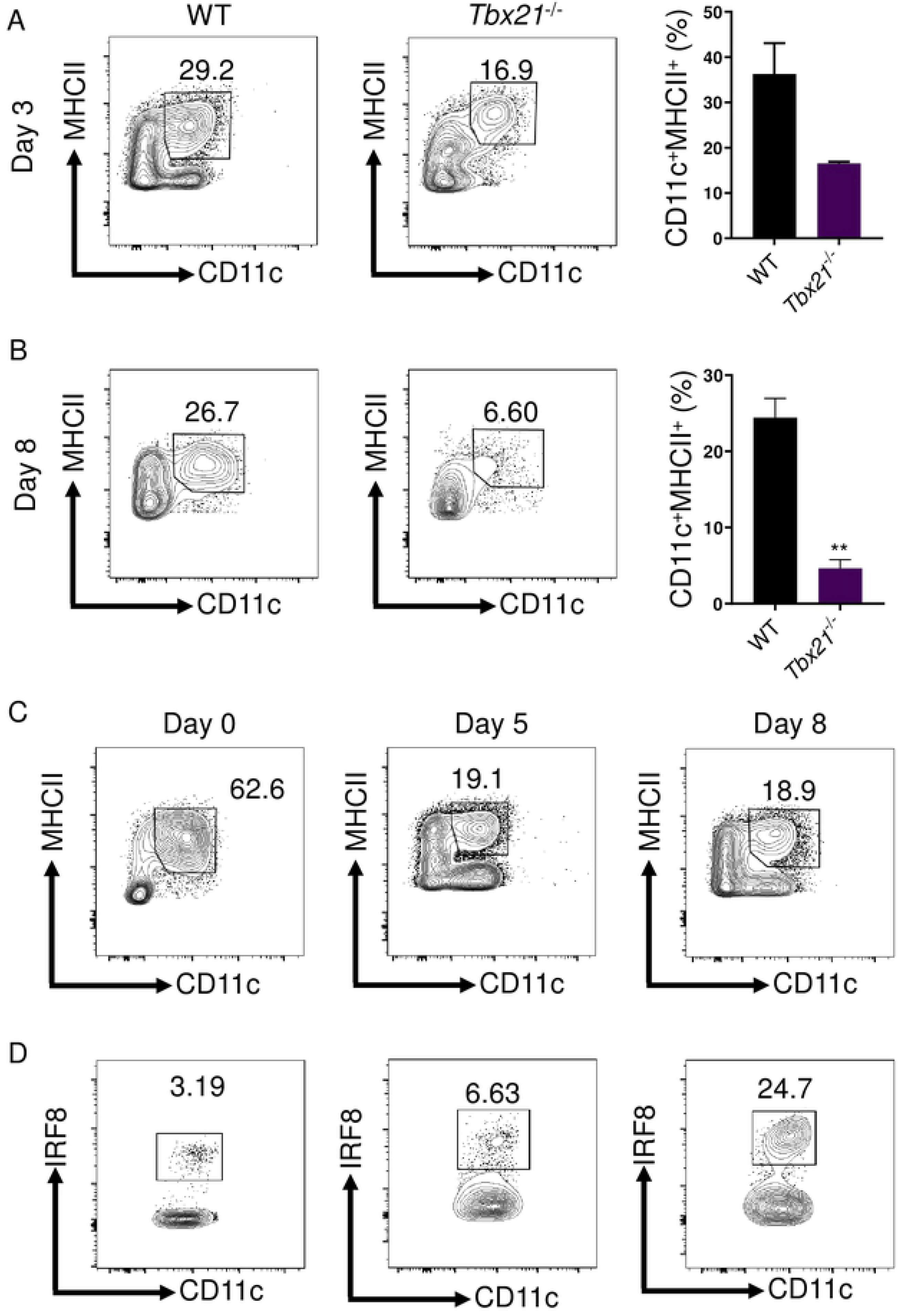
T-bet is required for maintaining inflammatory DCs during *T. gondii* infections. (**A-D**) WT, *RAG2*^−/−^, and *Tbx21*^−/−^ mice were infected i.p. with 20 cysts of *T*. *gondii*. (A, B) WT and *Tbx21*^−/−^ PECs were harvested on days 3 and 8 and Lin-CD11c+MHCII+ DCs and were analyzed by flow cytometry. (A, B) Average frequencies of Lin-CD11c+MHCII+ DCs in the PECs were analyzed on days 3 and 8 following infection. (C, D) *RAG2*^−/−^ PECs were harvested on days 0, 5, and 8 and Lin-CD11c+MHCII+ DCs and were analyzed by flow cytometry. (D) Average frequencies of Lin-CD11c+MHCII+IRF8+ DCs in PECs of *RAG2*^−/−^ mice were analyzed on days 0, 5, and 8 following infection. Results are representative of three-independent experiments involving at least 3 mice per group. Statistical analyses were done using unpaired t-test analysis of individual groups, \*\**P*<0.01. Error bars, standard error mean.

**Figure S2.**
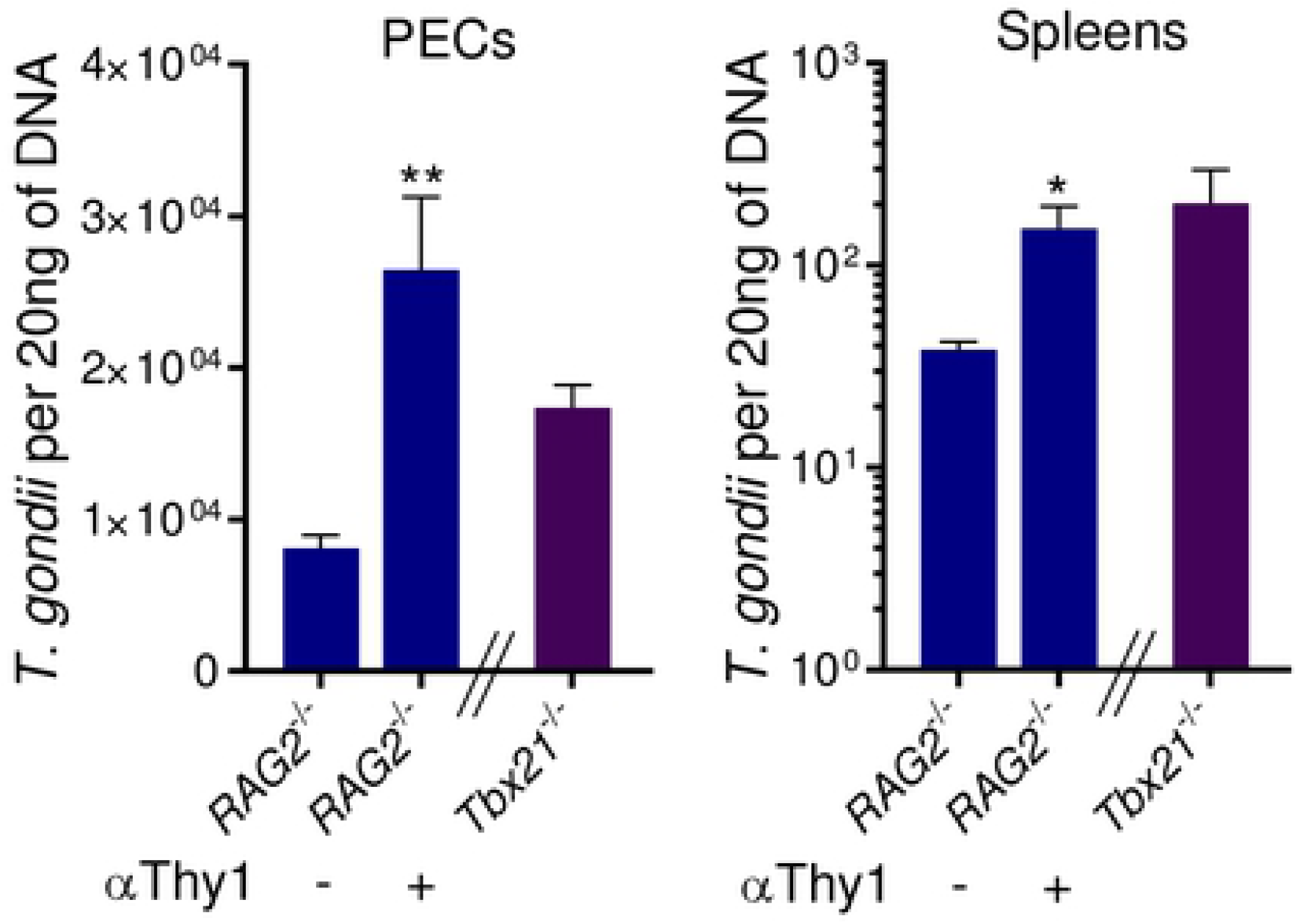
T-bet-dependent ILC1s are critical for parasite restriction. Parasite burden was assessed from PECs and spleen of *RAG2*^−/−^ mice treated with or without anti-Thy1 antibody by qPCR. Results are representative of three-independent experiments involving at least 3 mice per group. Error bars, standard error mean. Statistical analyses were done using unpaired t-test analysis of individual groups, \**P*<0.05, \*\**P*<0.01. Error bars, standard error mean.

**Figure S3.**
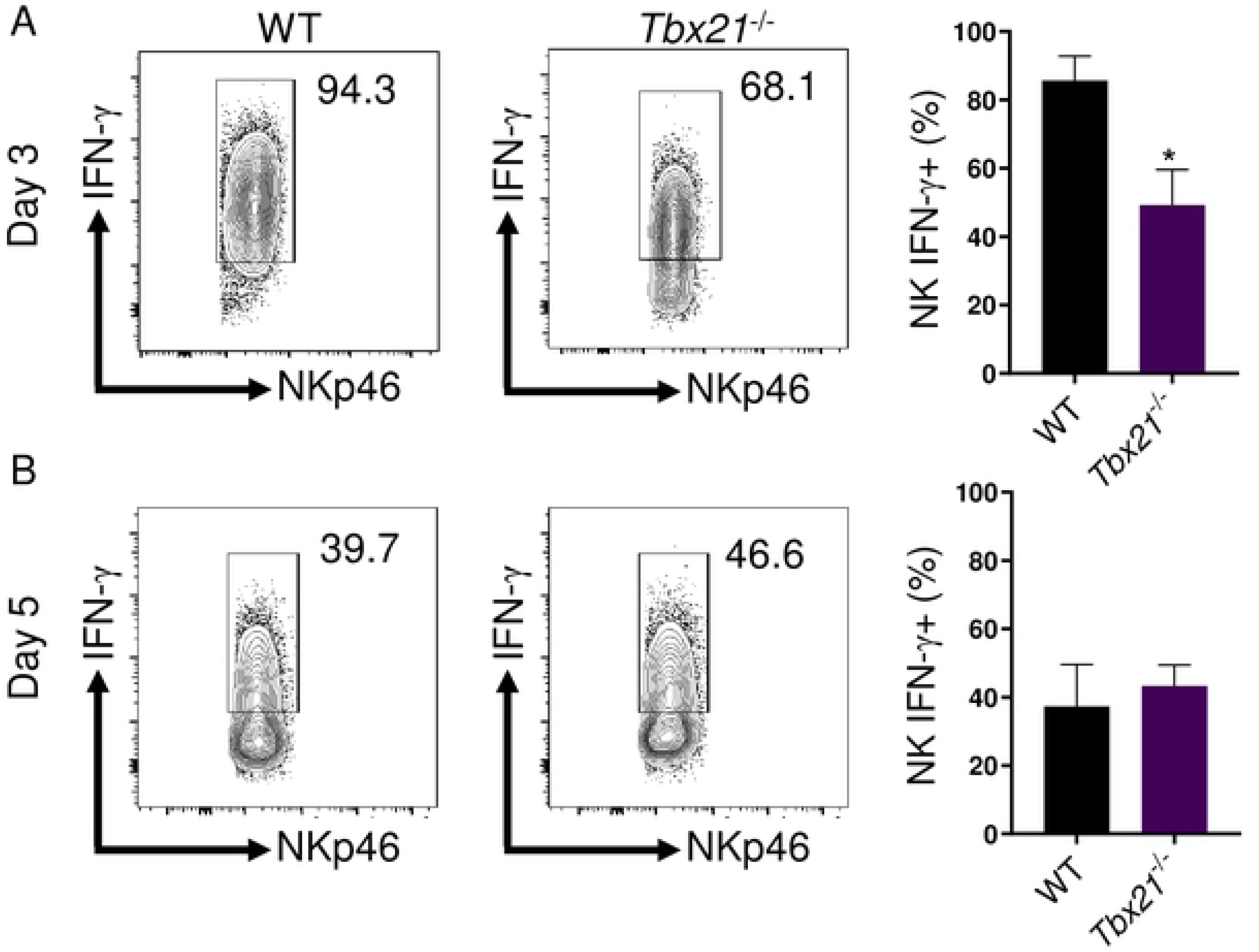
*T. gondii*-mediated NK-derived IFN-γ is T-bet-independent. (**A**, **B**) WT and *Tbx21*^−/−^ mice were infected i.p. with 20 cysts of *T*. *gondii*. Average frequencies of (A, B) CD127-NKp46+IFN-γ+ NK cells in the PECs were analyzed on days 3 and 5 following infection. Results are representative of three-independent experiments involving at least 3 mice per group. Statistical analyses were done using unpaired t-test analysis of individual groups, \**P*<0.05. Error bars, standard error mean.

**Figure S4.**
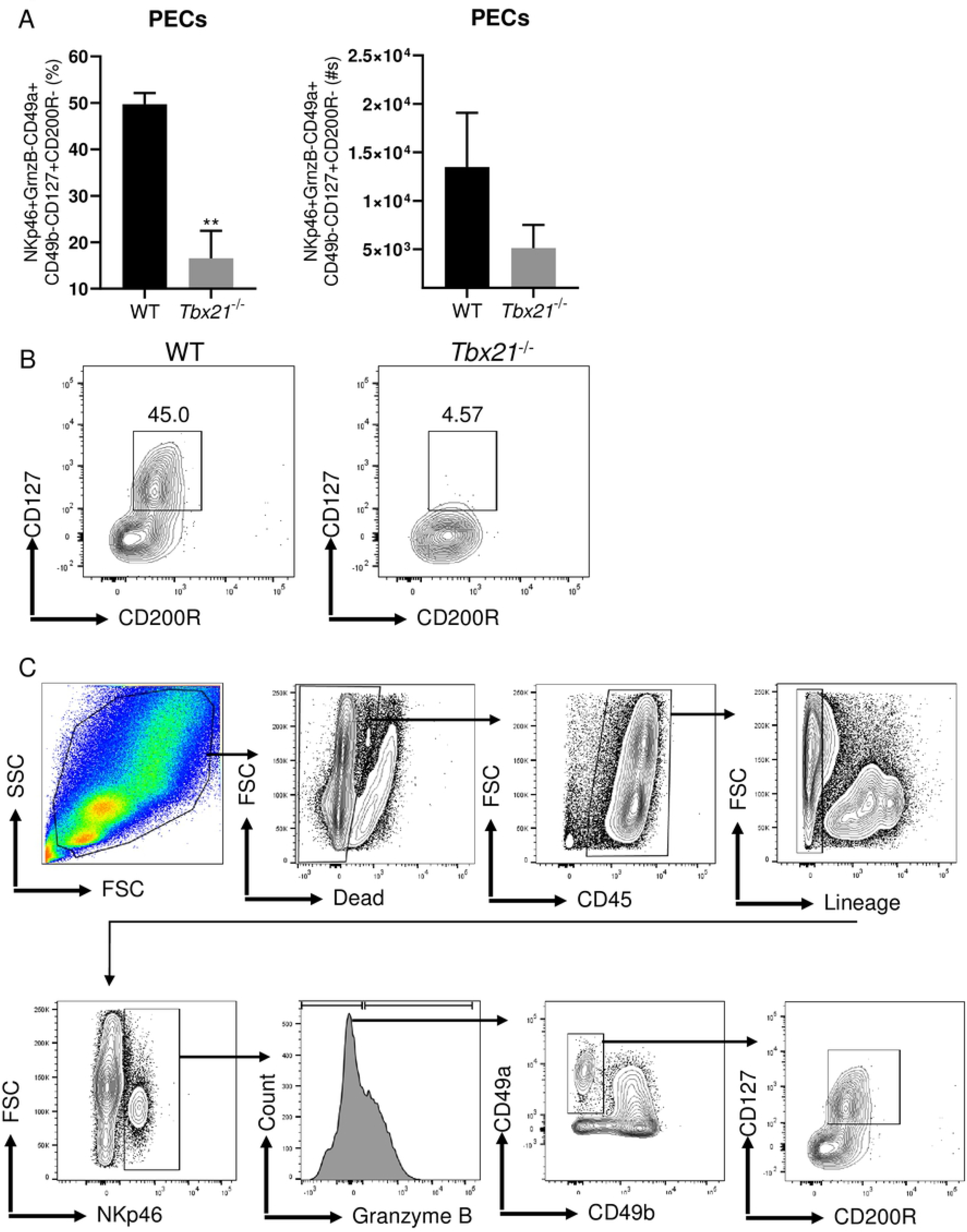
ILC1s are absent from *Tbx21*^−/−^ mice during *T. gondii infection*. (**A**, **B, C**) WT and *Tbx21*^−/−^ mice were infected i.p. with 20 cysts of *T*. *gondii* and PECs were assessed for ILC1s. (A, B) Average frequencies of CD45^+^CD3^−^CD19^−^Ly6G^−^NKp46^+^GranzymeB^−^CD49b^−^ CD49a^+^CD127^+^CD200R^+^ in the PECs were analyzed on day 5 following infection. (C) Representative gating strategy for ILC1s. Statistical analyses were done using unpaired t-test analysis of individual groups, \*\**P*<0.01. Error bars, standard error mean.

**Figure S5.**
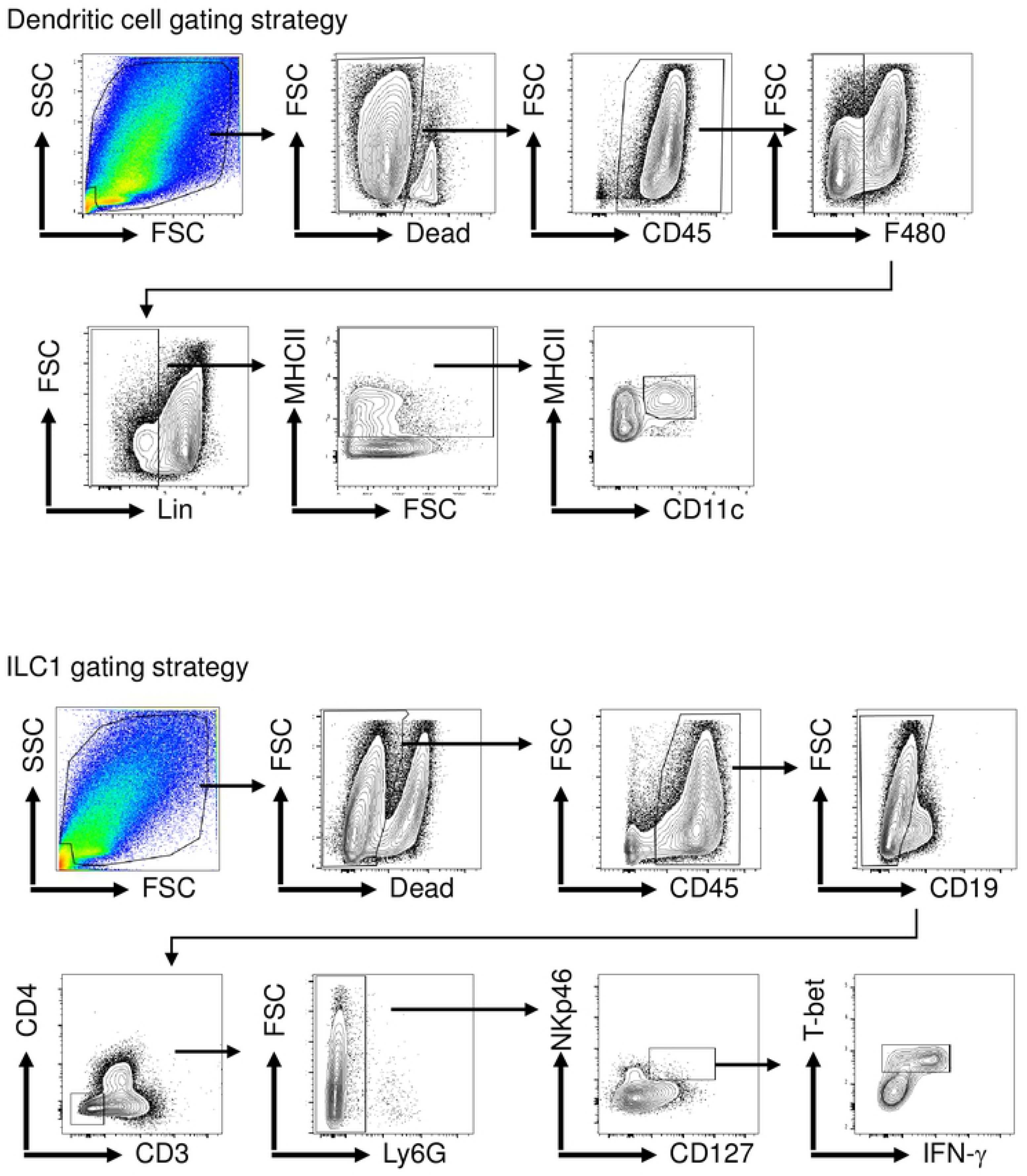
Representative gating strategies for DC and ILC1.

